# POM121 regulates lentiviral nuclear import in a capsid-host factor-dependent manner

**DOI:** 10.64898/2026.11.09.765341

**Authors:** Laura Arribas, Andrea Martínez-Ramas, James A. Cain, Steven Lynham, Michael H. Malim, Gilberto Betancor

## Abstract

Lentiviruses such as human immunodeficiency virus 1 (HIV-1) access the nuclear lumen through the nuclear pore complex (NPC), a process coordinated by the interactions between the viral capsid and nucleoporins (NUPs). Here, using unbiased proteomic screens, we identified several capsid-associated NUPs, including POM121. Depletion of POM121 impaired HIV-1 infection in multiple cell lines and in primary CD4+ T cells, whereas equine infectious anaemia virus (EIAV) infection was enhanced, indicating that POM121 effect is virus-specific. A cytoplasmic carboxy-terminal fragment of POM121, containing its FG-rich region inhibited HIV-1 and a broader range of lentiviruses. Analysis of HIV-1 capsid mutants showed that physical interaction with POM121 does not predict functional dependence, as capsids retaining POM121 binding remained insensitive to its depletion. Strikingly, loss of cyclophilin A (CypA) completely abolished HIV-1 dependence on POM121, whereas depletion of CPSF6 produced only partial rescue, demonstrating that POM121 sensitivity is dictated by the broader network of capsid-host interactions. Finally, POM121 depletion impaired HIV-1 nuclear accumulation and selectively disrupted the nuclear import of other viral and cellular cargo. Together, these findings identify POM121 as a selective regulator of lentiviral nuclear entry whose functional contribution is determined by capsid and host-factor context.

## Introduction

A defining feature of lentiviruses, such as human immunodeficiency virus 1 (HIV-1), is their ability to infect non-dividing cells. This capacity relies on their ability to exploit the nuclear pore complex (NPC) as a gate to gain access to the cell nucleus, where the proviral DNA integrates into the host chromosome [1].

The HIV-1 Capsid (CA) protein assembles into a lattice comprising approximately 250 CA hexamers and 12 pentamers arranged in the shape of a fullerene cone [2, 3]. This protein shell houses the viral genomic RNA together with the enzymes reverse transcriptase (RT) and integrase (IN), as well as additional viral and cellular components [4]. Initially considered a mere vessel that disassembled in the cell cytoplasm [5], the viral capsid is now recognized as a major hub for interactions with cellular proteins [5–7]. Mounting evidence shows that most of these interactions require an assembled capsid lattice, indicating that capsid disassembly must be tightly coordinated with the successive steps of early infection. This process, termed uncoating, remains the subject of intense investigation, with current models proposing that substantial capsid remodelling or disassembly occurs during passage through the NPC or after entry into the nuclear lumen [8–10]. Consistent with this view, recent studies employing nuclear pore-like structures reconstitution systems [11–13], and advanced microscopy approaches [10, 14] have demonstrated that intact, or nearly intact, capsids can reach the nuclear compartment.

As the capsid moves from the periphery of the cell membrane towards the nucleus, it sequentially engages different cellular factors. One of the earliest is cyclophilin A (CypA), which binds to the eponymous CypA-binding loop located around positions 85-93 of CA [15]. This interaction influences the subsequent interplay between capsid and host factors, including protection from human tripartite-containing motif 5α (TRIM5α)-mediated restriction [16]. Conversely, CA is also targeted by antiviral proteins, most notably the interferon-stimulated factors TRIM5α and myxovirus resistance protein 2 (MX2) [17–19].

At the nuclear envelope, the capsid engages components of the NPC. The NPC is a macromolecular structure composed of some 30 different nuclear pore proteins (most of them referred to as NUPs), present in multiple copies and organized into cytoplasmic filaments, a central scaffold and transport channel, and a nuclear basket extending into the nucleoplasm [20]. A subset of NUPs is enriched in phenylalanine-glycine (FG) repeats. These repeats occur in different canonical forms, including FG, GLFG, and FXFG motifs [21], and several FG-NUPs directly engage the conserved FG-binding pocket formed at the interface between adjacent CA subunits within the capsid lattice. These include cytoplasmic NUP358 [22], central channel-associated NUPs such as NUP62 or NUP98 [21, 23], and the nuclear basket component NUP153 [24, 25]. Once the capsid reaches the nuclear side of the pore, the nuclear protein CPSF6 also engages this pocket and contributes to capsid release from the NPC and subsequent trafficking towards transcriptionally active regions of the genome, thereby influencing integration-site selection [26, 27].

POM121 is an integral membrane nucleoporin that contributes to anchoring the NPC to the nuclear envelope [28, 29]. It contains an amino (N)-terminal transmembrane region and an extended carboxy (C)-terminal region rich in FG repeats that projects toward the transport channel [29]. Previous studies have implicated POM121 in HIV-1 nuclear entry: its depletion reduces HIV-1 infection and the nuclear accumulation of viral DNA without substantially affecting reverse transcription, while its FG-rich C-terminal contributes to its proviral activity [30, 31]. POM121 has also been shown to associate with assembled HIV-1 capsids through its C-terminal region, and its requirement during infection is influenced by CypA and CPSF6 (31). However, what determines whether a lentiviral capsid functionally depends on POM121, whether this dependence can be predicted from physical capsid binding, and how it is shaped by other capsid-associated host factors remain poorly understood.

Here, we investigate the determinants of POM121 dependency during lentiviral infection. POM121 was independently identified as a capsid-associated protein in an unbiased proteomic screen, and its depletion reduced HIV-1 infection in multiple cell lines and, importantly, in primary human CD4+ T cells. However, functional dependence on POM121 did not correlate with capsid binding. HIV-1 variants bearing substitutions in the CypA-binding loop, FG-binding pocket or tri-hexamer interface were resistant to POM121 depletion, despite several retaining the ability to bind POM121. Moreover, depletion of CypA completely abolished HIV-1 dependence on POM121, whereas CPSF6 loss produced only partial rescue, indicating that POM121 sensitivity is dictated by the broader network of capsid-host factor interactions.

We further show that POM121 dependency differs markedly among lentiviruses. Whereas SIVmac displayed a requirement for POM121, EIAV infection was enhanced in its absence, indicating that the same nucleoporin can exert opposing effects on different lentiviruses. Conversely, a cytoplasmic C-terminal fragment of POM121 inhibited a broader range of lentiviruses, demonstrating that the antiviral activity of this truncated form and dependence on endogenous POM121 are distinct properties. Finally, POM121 depletion impaired HIV-1 nuclear accumulation and selectively disrupted the nuclear import of viral and cellular nuclear localisation signal (NLS)-bearing cargos. Together, these findings establish POM121 as a selective regulator of nucleocytoplasmic transport and show that its contribution to lentiviral infection is determined by capsid sequence and by the host-factor network that engages with the capsid during nuclear entry.

## RESULTS

### Proteomic analysis of HIV-1 capsid interactors

To identify cellular proteins associated with the HIV-1 capsid, we performed an unbiased proteomic screen using capsid-nucleocapsid (CANC) assemblies, which have been demonstrated to recapitulate the interactions of the viral capsid [19, 24, 32]. Because changes in CA sequence can alter host factor engagement [6], we analysed in parallel CANC assemblies bearing the P90A substitution. This mutation disrupts CypA binding [33], confers resistance to MX2-mediated restriction [18, 34], and alters the dependence of HIV-1 on specific NUPs [35, 36]. MLV CANC assemblies and CANC assembly buffer were included as controls.

We performed stable isotope labelling by amino acids (SILAC)-based screens using U87-MG cell lysates under these four conditions. Following isolation of the CANC assemblies by centrifugation, associated cellular proteins were identified by LC-MS/MS analysis. As expected, well established capsid interactors, including CypA, CPSF6, NUP153, and NUP358/RANBP2 were enriched in wild type HIV-1 CANC pull-downs, whereas CypA was absent from P90A CANC samples (Fig 1A, S1 Fig). Interestingly, several NPC components were consistently enriched in HIV-1 CANC samples, including NUPL1/NUP58, NUP54, NUP160, NUP205, and POM121.

**Fig 1.**
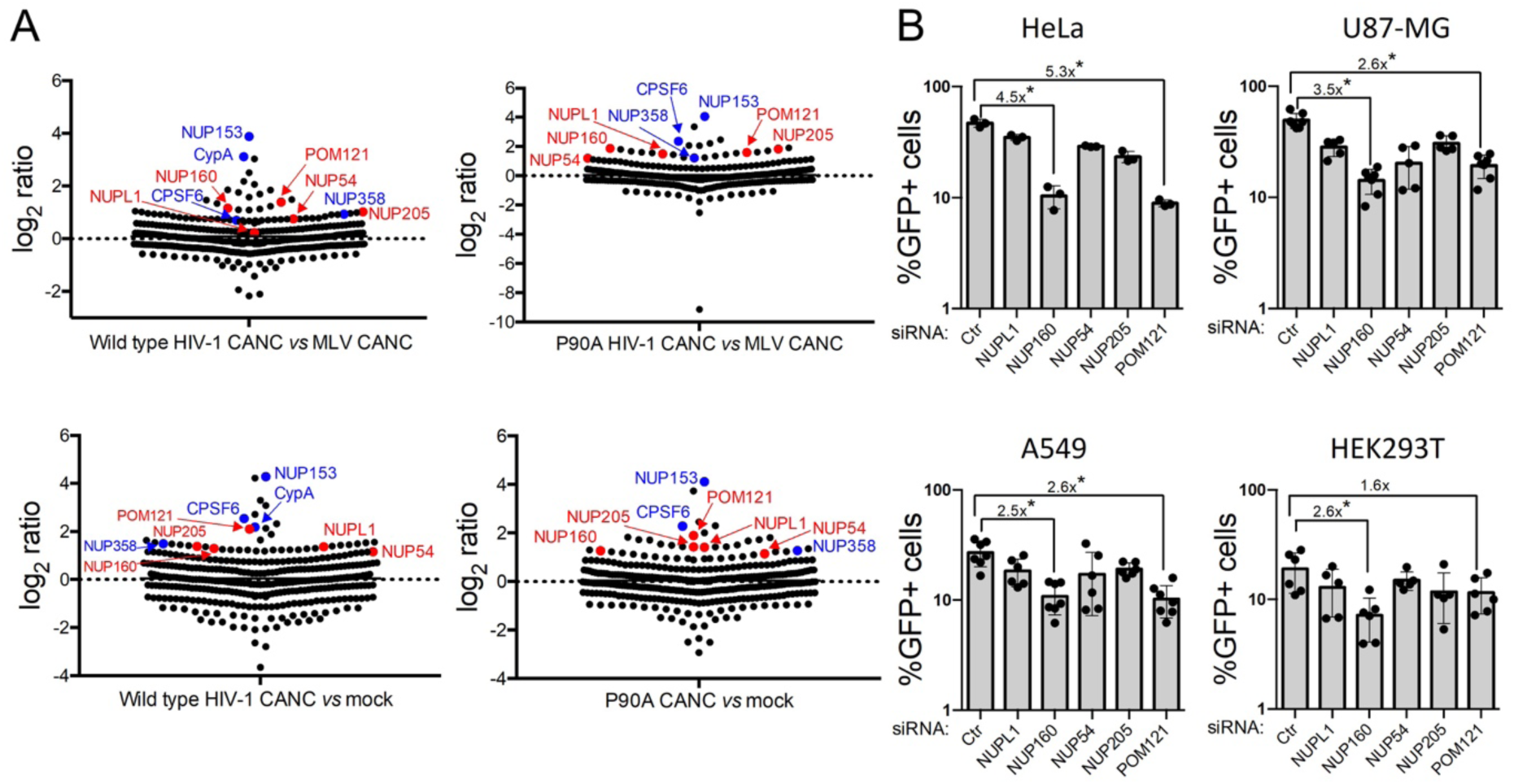
HIV-1 capsid interacts with multiple NUPs. (**A**) SILAC screen for capsid-binding proteins. Wild type and mutant P90A HIV-1 CANC assemblies, MLV CANC assemblies or CANC assembly buffer (mock) were incubated with HEK293T lysates from cells labelled with ‘light’, ‘medium’ or ‘heavy’ amino acids. Mixtures were subjected to centrifugation through a 70% sucrose cushion, washed with wash buffer, centrifuged again, and analysed by tandem mass spectrometry. Representative experiments comparing wild type CANCs (light) versus MLV CANCs (heavy) (n = 2 biological replicates), P90A CANCs (medium) versus MLV CANCs (heavy) (n = 2), wild type CANCs (medium) versus mock (heavy) (n = 2) and P90A CANCs (light) versus mock (heavy) (n = 2) are shown. Known HIV-1 capsid interactors are shown in blue, while the subset of NUPs identified in this study are shown in red. (**B**) HeLa, U87-MG, A549, and HEK293T cells were transfected twice, 24 h apart, with 20 nM of specific siRNAs (a non-targeting siRNA was included as a control, Ctr). 24 h later, cells were challenged with a HIV-1 based lentiviral vector expressing GFP (HIV-1/GFP) and the percentage of infected (GFP-expressing) cells quantitated 48 h later by flow cytometry. The fold reduction in viral infectivity between cells transfected with Ctr siRNA and those transfected with siRNAs against *NUP160* and *POM121* is shown (n = at least 3; mean ± standard deviation (SD); * p-value < 0.05; unpaired t-test to Ctr siRNA).

We next investigated whether these candidate NUPs contributed functionally to HIV-1 infection. Because NUP expression and functional requirements can vary across cellular backgrounds [37], we performed the analysis in four different cell lines. U87-MG, HeLa, A549 and HEK293T cells were transfected twice with siRNAs directed against these NUPs, or with a control siRNA of scrambled sequence, and were subsequently challenged with an HIV-1-based reporter vector encoding GFP (HIV-1/GFP). Depletion of NUPL1, NUP54, and NUP205 did not detectably affect HIV-1 infectivity under these conditions (Fig 1B). In contrast, viral infectivity was consistently reduced in cells depleted of NUP160 and POM121, although the magnitude of the phenotype varied between cellular backgrounds.

To examine whether these effects reflected a general perturbation of viral infection, we challenged the same cells with murine leukaemia virus (MLV), a gammaretrovirus dependent on nuclear envelope rupture during mitosis to access the cell nucleus. Depletion of POM121 or NUP160 did not impair MLV infection (S2 Fig), supporting a specific role for these proteins in HIV-1 replication.

### POM121 depletion differently affects infection by HIV-1 capsid mutants and other lentiviruses

Amongst the nucleoporins identified in our screen, we focused on POM121 because of its direct association with HIV-1 CANC assemblies, its position as an FG-containing transmembrane component of the NPC, and previous studies implicating POM121 in HIV-1 nuclear entry and capsid-dependent infection [30, 31]. To investigate its contribution to viral infection, we generated a bulk population and two clonal HeLa cell lines in which the endogenous *POM121* alleles were suppressed by clustered regularly interspaced short palindromic repeats (CRISPR)-associated protein 9 (Cas9) genome editing, together with a control CRISPR line [38] Challenge of these cells with HIV-1/GFP confirmed the phenotype observed following siRNA-mediated depletion, with infection significantly reduced in the bulk population and, more prominently, in the clonal POM121 CRISPR lines (Fig 2A). A similar reduction was observed using replication-competent full-length HIV-1 (Fig 2B), indicating that the phenotype was not restricted to the lentiviral reporter system. To further examine the relationship between POM121 and nuclear entry, cells were treated with aphidicolin to inhibit cellular DNA replication and cell-cycle progression, thereby preventing mitotic nuclear-envelope breakdown. Under these conditions, HIV-1 infection was further reduced in POM121 CRISPR cells (S3 Fig), consistent with a role for POM121 in NPC-dependent infection.

**Fig 2.**
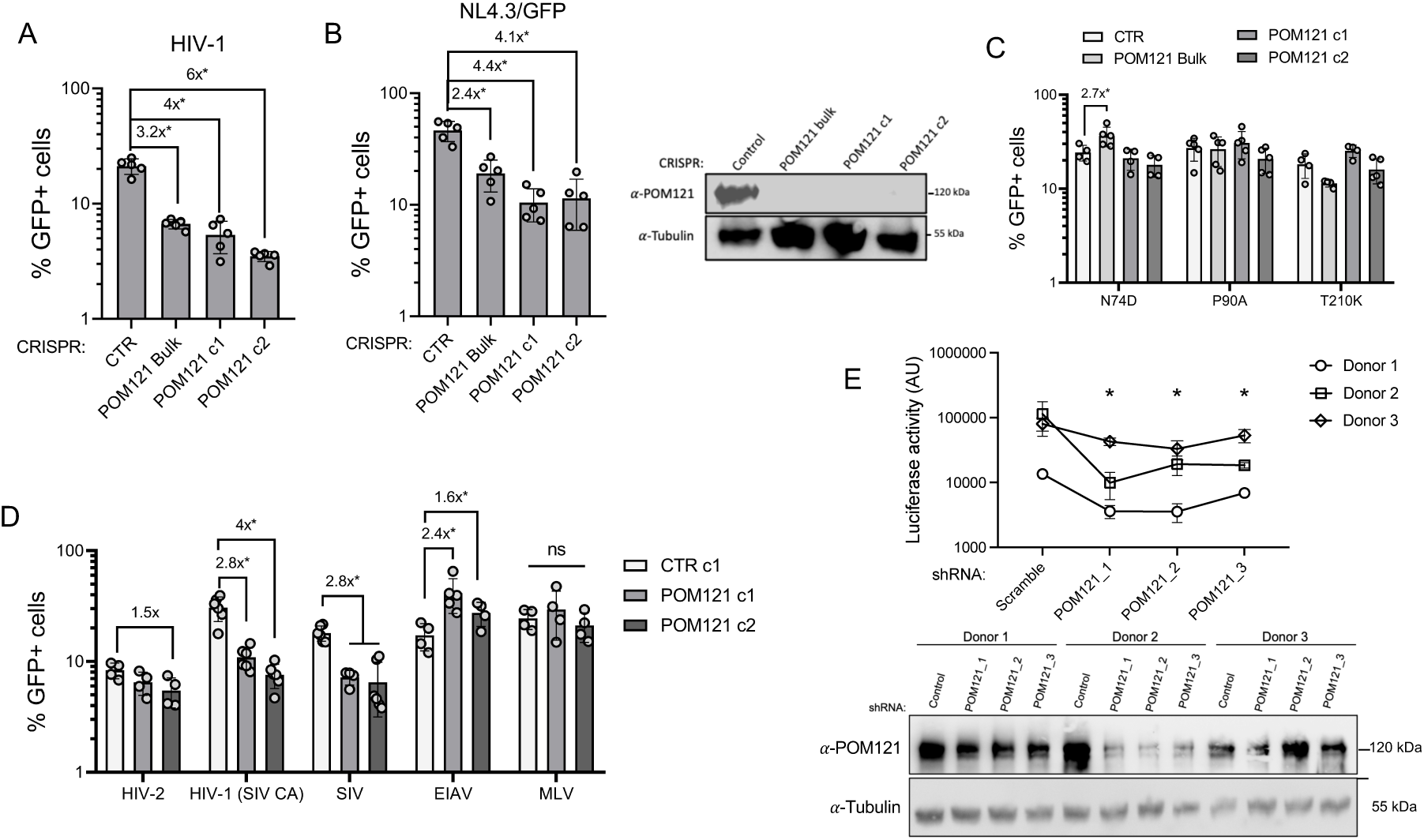
POM121 depletion inhibits HIV-1 in primary CD4 + T cells, but its effect in infection its capsid-dependent. (**A, B**) HeLa cells transduced with a control guide RNA (CTR), as well as a bulk population and two clonal cell lines (c1 and c2) in which *POM121* had been disrupted by CRISPR-Cas9 genome editing were challenged with HIV-1/GFP (**A**) or with NL4.3/Nef-IRES-GFP (**B**) and infection was quantified at 48 h by flow cytometry (n = 5 biological replicates; mean ± SD; * p-value < 0.05; unpaired t-test to CTR CRISPR cells). The right panel shows the immunoblot analysis of HeLa CRISPR cells shown in (**A, B**). (**C**) same as (**A, B**), but cells were challenged with HIV-1 capsid mutants N74D, P90A, T210K (n = at least 3 biological replicates; mean ± SD; * p-value < 0.05; unpaired t-test to CTR CRISPR cells). (**D**) HeLa CTR CRISPR cells and CRISPR POM121 clonal lines were challenged with lentiviral particles based on HIV-2, HIV-1 (SIV CA), SIV, EIAV, and MLV, and the number of GFP-expressing cells measured by flow cytometry 48 h later (n = 4 biological replicates; mean ± SD; * p-value < 0.05; unpaired t-test to CTR CRISPR cells). (**E**) Primary CD4+ T cells were isolated from 3 donors, transduced with shRNAs targeting *POM121* or a scrambled sequence shRNA and challenged with NL4.3/Nef-IRES-Renilla. The luciferase activity was determined 48 h later. Up: raw infectivity data from all three donors (n = 3 technical replicates; mean ± SD; * p-value < 0.05; unpaired t-test to scramble shRNA); down: immunoblot analysis of POM121 expression in scramble- and *POM121*-shRNA transduced cells.

The HIV-1 capsid harbours different interacting surfaces, including the CypA-binding loop [15], the FG-binding pocket [7], and the tri-hexamer interface [25, 39]. To determine whether these surfaces influence POM121 dependency, we challenged our CRISPR cell lines with viruses bearing P90A, N74D, or T210K CA substitutions, affecting these three regions, respectively. None of these mutants were impaired by POM121 depletion (Fig 2C). Thus, alterations at structurally distinct capsid interfaces are sufficient to uncouple HIV-1 infection from its dependence on POM121. Notably, N74D and P90A have also been previously reported to confer resistance to MX2-mediated restriction [18, 34].

We next asked whether POM121 dependency was conserved among lentiviruses. Lentiviruses differ substantially in the sequence and functional use of capsid surfaces engaged by host factors. CypA interactions vary amongst HIV-2, SIVmac (simian immunodeficiency virus derived from macaque) and EIAV (equine infectious anaemia virus) [40, 41], whereas conservation and functional usage of the FG-binding pocket also differ amongst lentiviruses [42]. The higher-order architecture of the tri-hexamer interface is more broadly conserved, although the precise host-factor interactions mediated by this surface remain less well defined outside HIV-1 [6]. We therefore challenged the POM121 CRISPR clones with HIV-2, SIVmac, and EIAV, as well as a chimeric HIV-1 carrying SIVmac CA in an otherwise HIV-1 background [HIV-1(SIV CA)] (Fig 2D). Interestingly, most lentiviruses tested were refractory to POM121 depletion. In contrast, viruses bearing SIVmac CA, both SIVmac itself and HIV-1(SIV CA), showed an ∼2-3-fold reduction in infectivity. Remarkably, EIAV displayed the opposite phenotype, with infection increasing ∼2-3-fold in POM121 CRISPR cells relative to control cells. These results indicate that the effect of POM121 on lentiviral infection can range from proviral to inhibitory depending on the viral capsid context.

### POM121 depletion reduces HIV-1 infectivity in primary CD4+ T cells

Because NPC composition and nucleoporin requirements can vary between cellular backgrounds [35, 37], we sought to determine whether the POM121 phenotype observed in cell lines was also maintained in primary human CD4+ T cells, the main target of HIV-1 infection *in vivo*. Cells from three healthy donors were transduced with short hairpin RNAs (shRNAs) specific for *POM121*, as well as a negative control (scrambled sequence) and subsequently challenged with the NL4.3 virus.

POM121 depletion significantly reduced HIV-1 infection in cells from all three donors (Fig 2E). The magnitude of the antiviral phenotype paralleled the extent of POM121 depletion detected by immunoblot, with donors one and two showing both the strongest reduction in POM121 expression and the largest decrease in infectivity, whereas donor three displayed a more modest effect. These results extend the requirement for POM121 beyond transformed cell lines and establish its contribution to efficient HIV-1 infection in primary human CD4+ T cells.

### The carboxy-terminal portion of POM121 inhibits lentiviral infection

Previous studies have demonstrated that overexpression of N-terminally truncated POM121 proteins inhibits HIV-1 infection [31, 43, 44]. We interrogated this notion and split the canonical 1,249-residue protein into two fragments comprising residues 1-682 (POM121-ΔC) and residues 683-1249 (POM121-ΔN) (Fig 3A). Full-length POM121, both truncated proteins, and MX1 as a negative control were stably expressed in HeLa cells by lentiviral transduction (Fig 3B). We next examined the cellular localisation of the truncate proteins. Consistent with a recent report [44], POM121-ΔN, which lacks the transmembrane domain, accumulates in the cell cytoplasm, while POM121-ΔC showed a distribution similar to the full-length protein, and predominantly localises at the nuclear envelope (Fig 3C).

**Fig 3.**
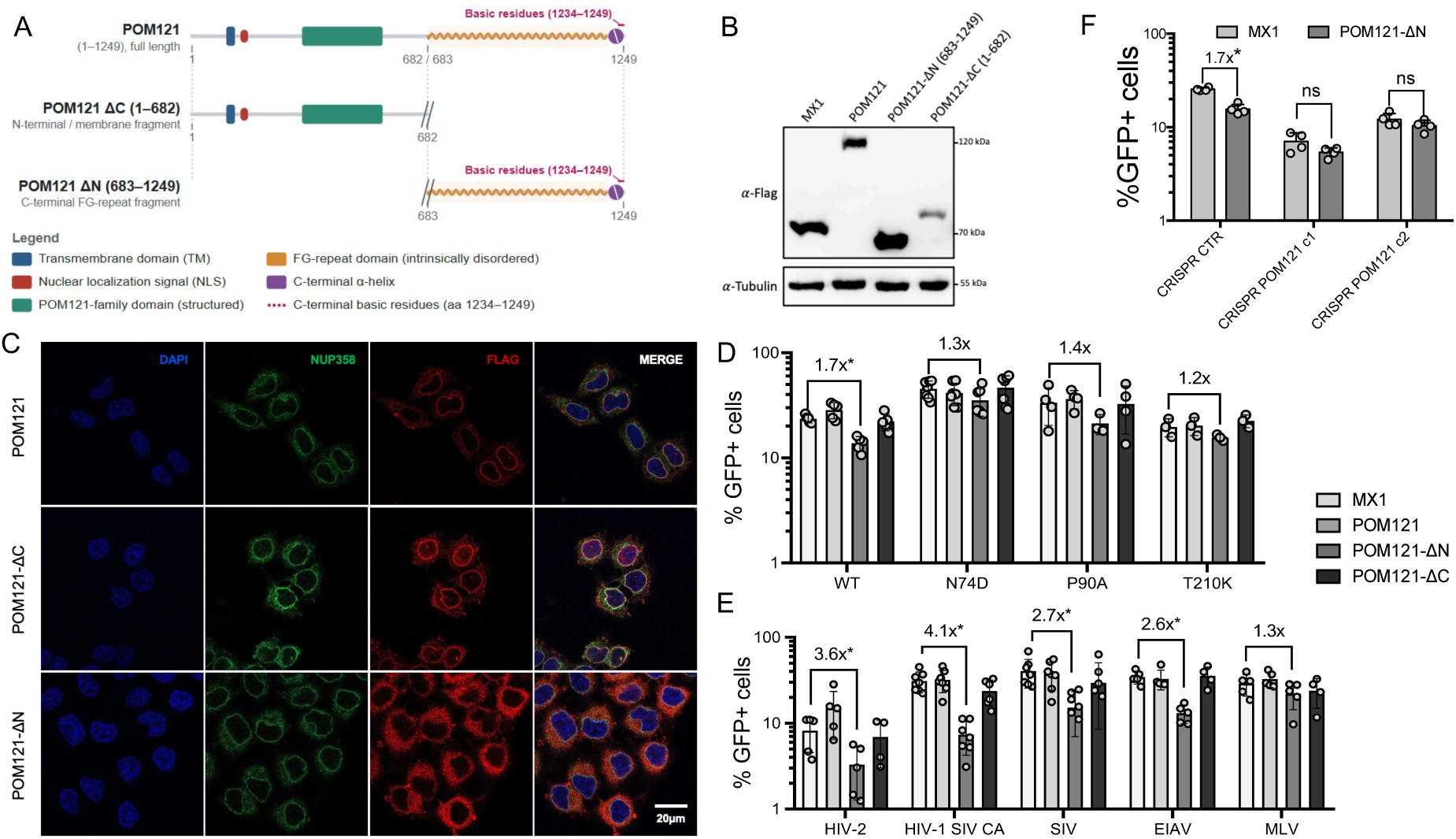
A carboxy-terminal fragment of POM121 inhibits lentiviral infection. (**A**) Schematic representation of canonical POM121 and truncated forms POM121 1-682 (POM121-ΔC) and POM121 683-1249 (POM121-ΔN). The main protein structural and functional domains are shown. (**B**) Immunoblot analysis of canonical and truncated POM121 forms expression. (C) HeLa cells were stably transduced with puromycinR lentiviral vectors encoding Flag-tagged POM121, POM121-ΔN, and POM121-ΔC. Cells were seeded onto glass coverslips and localization of POM121 forms and endogenous NUP358 were determined by immunostaining with anti-Flag and anti-NUP358 antibodies and confocal microscopy. DAPI was used to stain the nuclei. (**D**) HeLa cells were stably transduced with lentiviral particles expressing MX1 (negative control), full-length POM121, POM121-ΔN, and POM121-ΔC. Cells were challenged with wild type and mutants N74D, P90A, and T210K HIV-1/GFP and infection was quantified at 48 h by flow cytometry (n = at least 3 biological replicates; mean ± SD; * p-value < 0.05; unpaired t-test to MX1 cells). (**E**) sames as (**D**), but cells were challenged with HIV-2, HIV-1 (SIV CA), SIV, EIAV, and MLV particles (n = at least 4 biological replicates; mean ± SD; * p-value < 0.05; unpaired t-test to MX1 cells). (**F**) Control CRISPR cells and clonal POM121 CRISPR populations 1 and 2 were stable transfected with puromycinR particles encoding MX1 or POM121-ΔN. Cells were then challenged with HIV-1/GFP and infection was quantified at 48 h by flow cytometry (n = 4 biological replicates; mean ± SD; * p-value < 0.05; unpaired t-test to MX1 cells).

HIV-1 infection was modestly but significantly impaired in cells expressing POM121-ΔN, while it remained unaffected by expression of the full-length or ΔC proteins (Fig 3D). Thus, the C-terminal POM121-ΔN fragment, which retains the FG-rich region, can inhibit HIV-1 infection. We next asked whether this inhibitory activity extended to capsid mutants and other lentiviruses. POM121-ΔN slightly reduced infection by the P90A mutant, whereas N74D and T210K remained resistant (Fig 3D). Surprisingly, all other lentiviruses tested were inhibited by POM121-ΔN, with HIV-2, SIVmac, and EIAV showing an ∼2-3-fold reduction in infectivity and the chimeric HIV-1(SIV CA) virus displaying the strongest phenotype, at ∼4-fold (Fig 3E). As expected, MLV infection was unaffected by POM121-ΔN expression. Notably, several lentiviruses that were not inhibited by depletion of endogenous POM121, including HIV-2 and EIAV, remained susceptible to inhibition by the isolated C-terminal region. These results indicate that sensitivity to the POM121-ΔN fragment is broader than functional dependence on endogenous POM121.

We finally asked whether inhibition by POM121-ΔN was additive to the infectivity defect caused by loss of endogenous POM121. MX1 or POM121-ΔN were therefore expressed in control and POM121 CRISPR HeLa cells. Importantly, the CRISPR guide employed to generate the POM121-deficient lines is directed to the N-terminal end of the protein, and therefore does not prevent the expression of the truncated protein. As expected, MX1 expression had no effect on infection, while POM121-deficient cells expressing MX1 showed the characteristic reduction in HIV-1 infectivity relative to control cells (Fig 3F). Interestingly, while overexpression of POM121-ΔN inhibited HIV-1 infection in control cells, no additional reduction was observed when the fragment was expressed in POM121-deficient cells. Therefore, the inhibitory effect of POM121-ΔN is non-additive with loss of endogenous POM121, consistent with both perturbations affecting the same or overlapping steps of HIV-1 infection.

### Sensitivity to POM121 depletion does not correlate with capsid binding

Our initial capsid-interactor screen identified POM121 as a binding partner of both wild type and P90A HIV-1 CANC assemblies. However, the P90A virus was insensitive to POM121 depletion. We therefore examined whether the other POM121-independent capsid mutants identified above, N74D and T210K, retained the ability to interact with POM121. Wild type, N74D, P90A, and T210K CANC complexes were incubated with whole-cell lysates from transfected 293T cells expressing MX1 as negative control, CPSF6 as positive control, POM121, POM121-ΔN and POM121-ΔC.

As expected, wild type CANC complexes readily precipitated CPSF6 and full-length POM121, confirming the results of our initial screen (Fig 4A). POM121-ΔN also interacted with wild-type CANC, whereas POM121-ΔC did not, consistent with previous reports implicating the C-terminal region of POM121 in capsid binding [31, 44]. P90A CANC assemblies displayed a similar interaction profile, precipitating CPSF6, full-length POM121 and POM121-ΔN (Fig 4B). T210K CANC complexes behaved similarly, retaining interaction with CPSF6, full-length POM121 and POM121-ΔN (Fig 4C).

**Fig 4.**
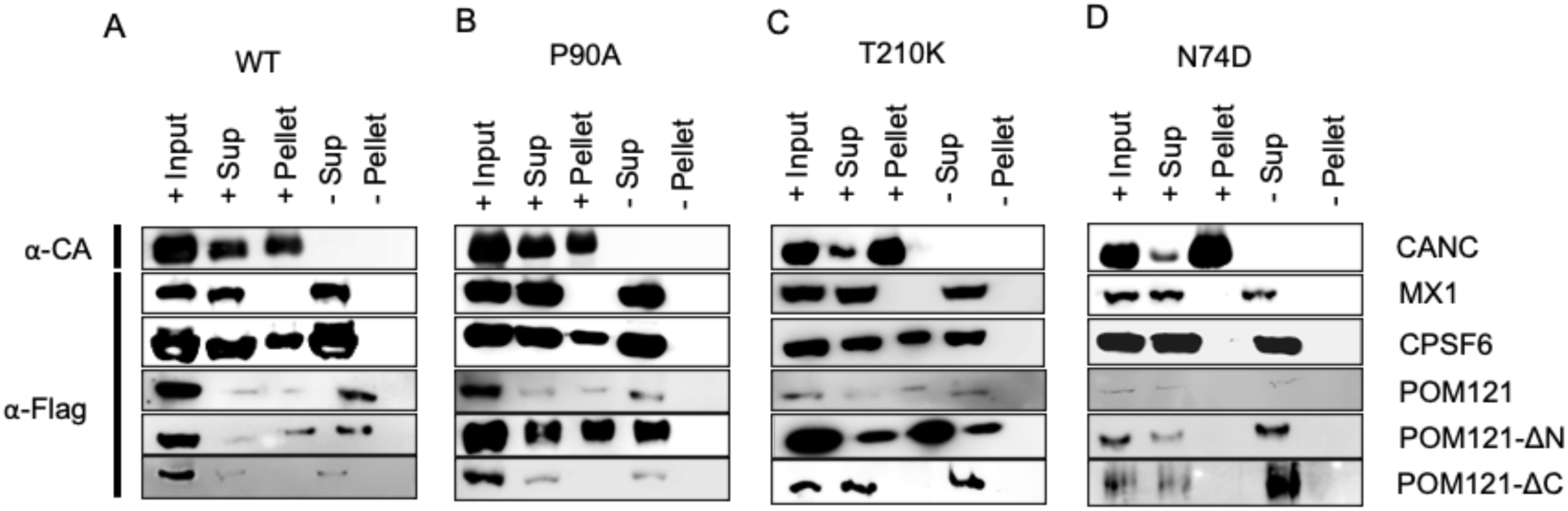
POM121 interaction with HIV-1 capsids. HEK293T cells were transfected with vectors expressing Flag-tagged MX1, CPSF6, POM121, POM121-ΔN, or POM121-ΔC. Cell lysates were incubated in the presence or absence of wild type (**A**), P90A (**B**), T210K (**C**), or N74D (D) CANC assemblies and subjected to centrifugation through a 70% sucrose cushion. Supernatant (sup), pellet and input samples were analysed by immunoblot using an anti-Flag antibody. A representative immunoblot for CANC is shown. All experiments were done at least 3 times.

In contrast, N74D CANC assemblies failed to precipitate CPSF6, as previously reported [45], and did not detectably interact with any of the POM121 constructs tested (Fig 4D). Thus, under our experimental conditions, the N74D substitution disrupts POM121 association with assembled CA. This differs from a previous report describing interaction between N74D CA assemblies and a C-terminal POM121 fragment [31].

Importantly, P90A and T210K provide a clear dissociation between POM121 binding and functional dependence: both capsids retained interaction with POM121, yet viruses bearing either substitution were resistant to POM121 depletion. Together, these results demonstrate that capsid association with POM121 is not sufficient to predict whether infection will depend on endogenous POM121.

### CypA and CPSF6 differentially contribute to POM121 sensitivity

Given that N74D and P90A capsid mutants were resistant to POM121 depletion, we asked whether the cellular factors whose binding is disrupted by these substitutions, CPSF6 and CypA, respectively, influence POM121 dependency. We therefore analysed HIV-1 infection in control, POM121, CypA and CPSF6 CRISPR cells, as well as in cells depleted simultaneously of POM121 and either CypA or CPSF6. Infection by HIV-1/GFP or full-length HIV-1 was not detectably affected by depletion of CypA or CPSF6 alone (Fig 5A, Fig 5B), in line with previous results [31, 45]. In contrast, loss of CypA in POM121 CRISPR cells fully restored wild type HIV-1 infectivity to the levels observed in control cells. CPSF6 depletion also increased HIV-1 infectivity in POM121-deficient cells, but the rescue was incomplete, with infection remaining approximately twofold lower than in control cells.

**Fig 5.**
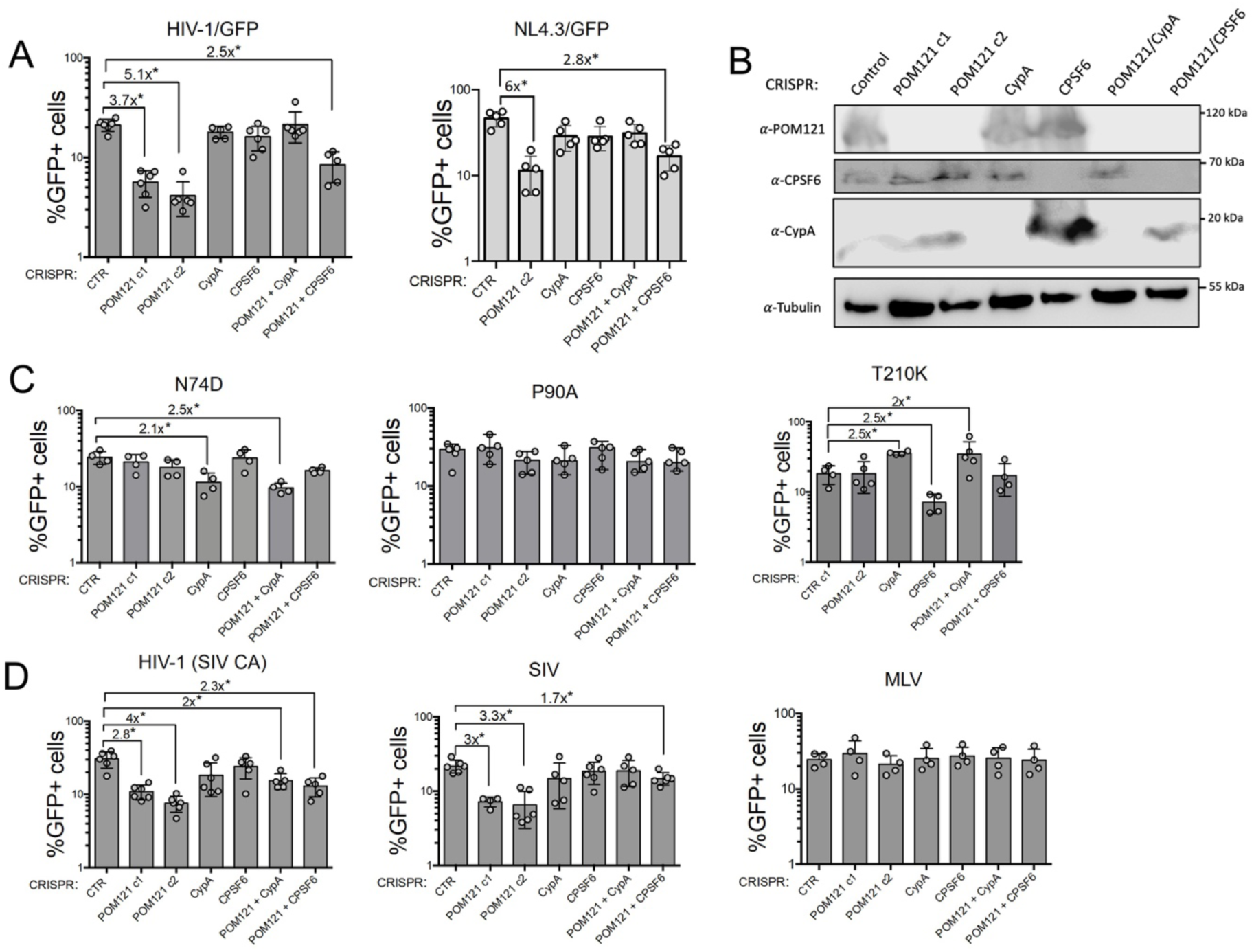
CypA is necessary for the functional dependence on POM121. (**A**) Parental HeLa cells and HeLa POM121 CRISPR cells were depleted of CypA or CPSF6 by CRISPR-Cas9 genome editing to generate CRISPR CypA, CRISPR CPSF6, CRISPR POM121 + CypA, and CRISPR POM121 + CPSF6 cells. These cells, together with CTR CRISPR cells, were challenged with HIV-1/GFP particles (left) or full-length NL4.3/Nef-IRES-GFP (right) and infectivity quantitated 48 h later by flow cytometry as the number of GFP+ cells (n = 5 biological replicates; mean ± SD; * p-value < 0.05; unpaired t-test to CTR cells). (**B**) Immunoblot analysis confirming depletion of targeted proteins. (**C**) Same as (**A**), but cells were challenged with capsid mutant viruses N74D, P90A, and T210K (n = at least 4 biological replicates; mean ± SD; * p-value < 0.05; unpaired t-test to CTR or double CTR cells). (**D**) Same as (**A**), but cells were challenged with HIV-1 (SIV CA), SIV, and MLV (n = at least 4 biological replicates; mean ± SD; * p-value < 0.05; unpaired t-test to CTR or double CTR cells).

We next examined the behaviour of the POM121-independent capsid mutants. N74D virus infection was reduced in CypA-deficient cells but was unaffected by CPSF6 depletion, as previously reported [36, 46]. The P90A virus was insensitive to loss of either CypA or CPSF6 (Fig 5C). Notably, T210K infection was strongly enhanced in the absence of CypA, increasing ∼4-fold, whereas CPSF6 depletion has no detectable effect.

We then extended this analysis to the other lentiviruses tested. HIV-2 and EIAV infection was unaffected by depletion of any of the proteins analysed here (S4 Fig). In contrast, SIVmac and the chimeric HIV-1(SIV CA) virus displayed a pattern similar to wild-type HIV-1: loss of CypA fully restored infectivity in POM121-deficient cells, whereas CPSF6 depletion produced only a partial rescue (Fig 5D).

Together, these results identify CypA as a major determinant of POM121 dependency in HIV-1 and SIVmac infection. CPSF6 also contributes to this phenotype, but its depletion is insufficient to fully bypass the requirement for POM121.

### POM121 regulates nuclear import of HIV-1 and other cargo

To get insight into the mechanism underlying the role of POM121 in HIV-1 infection, we first examined its cellular distribution. As expected, POM121 primarily localizes at the nuclear envelope in CTR CRISPR cells, whereas its signal is lost in POM121 CRISPR cell lines (S5 Fig). POM121 participates in NPC assembly and contributes to anchoring the pore complex at the nuclear envelope [28, 29]. However, its role in the nucleocytoplasmic transport of cargo remains ill-defined. We therefore used a system developed by Kane and colleagues [35], in which a chimeric GFP-LacZ reporter is fused to different nuclear localisation signals (NLSs), to examine the contribution of POM121 to nuclear transport. A panel of nine GFP-LacZ-NLS reporters was analysed in control and POM121 CRISPR HeLa cells. Nuclear accumulation of most reporters was unaffected by POM121 depletion, indicating that loss of POM121 does not cause a general defect in nuclear import (S6 Fig). Remarkably, however, we found that POM121 depletion strongly reduced the nuclear accumulation of GFP-LacZ constructs bearing the SV40 large T-antigen NLS or residues 1-25 of MX2 (N_MX2_) (Fig 6A, 6B). These results indicate that POM121 selectively contributes to the nuclear import of particular NLS-bearing cargos.

**Fig 6.**
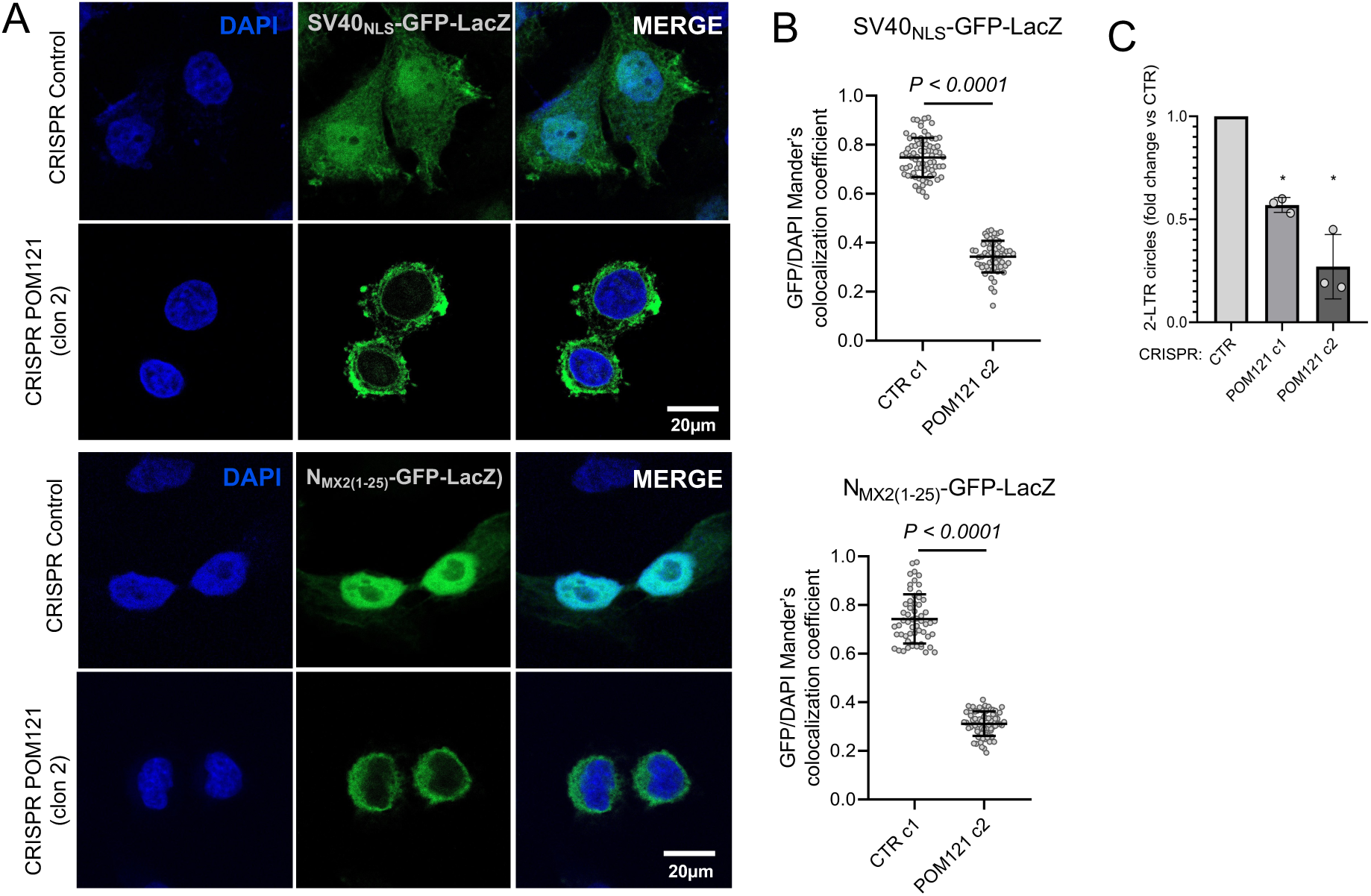
POM121 cellular distribution and effect on nuclear import. (**A**) HeLa control CRISPR and CRISPR POM121 cells stably expressing the SV40 large T antigen_NLS_-GFP-LacZ chimaera or the N_MX2(1-25)_-GFP-LacZ chimaera were seeded on glass coverslips. Cellular distribution of chimeric proteins was detected by confocal microscopy, with the nuclei stained with DAPI. (**B**) Nuclear localization of SV40 large T antigen NLS-GFP-LacZ and N_MX2(1-25)_-GFP-LacZ was determined for single cells by quantifying colocalization with DAPI using Mander’s coefficient for an average of 65 randomly selected cells (two-tailed unpaired t-test). (**C**) Quantification of HIV-1 2-LTR circles in control (CTR) and POM121 CRISPR cells (clones c1 and c2). Cells were infected with VSV-G-pseudotyped HIV-1 NL4.3/Nef-IRES-GFP, and total DNA was extracted 24 h later. 2-LTR circles were quantified by SYBR Green qPCR, normalised to the cellular reference gene *TRIM5* (ΔΔCt method) and expressed relative to CTR cells, set to 1 in each experiment. Bars show mean ± SD. * p < 0.005 (unpaired Student’s t-test) relative to CTR (n = 3).

We next examined whether POM121 depletion similarly affected HIV-1 nuclear entry. Accumulation of 2-LTR circles, commonly used as a surrogate marker of HIV-1 nuclear import, was significantly reduced in POM121 CRISPR cells (Fig 6C), consistent with previous studies [30, 31]. Thus, loss of POM121 impairs HIV-1 nuclear accumulation in addition to selectively affecting the transport of specific NLS-bearing cargos.

Finally, we examined whether HIV-1 infection altered the cellular distribution of full-length or truncated POM121 proteins. HeLa cells stably expressing full-length POM121, POM121-ΔN, or POM121-ΔC were infected with HIV-1/GFP and analysed by immunofluorescence 16 h later. As observed in uninfected cells, POM121-ΔN remained predominantly cytoplasmic, whereas POM121-ΔC displayed a distribution similar to full-length POM121 and accumulated at the nuclear envelope (S7 Fig). HIV-1 infection did not detectably alter the distribution of any of the POM121 constructs.

## DISCUSSION

The viral capsid plays a central role in coordinating trafficking from the cytoplasm to the nuclear compartment through interactions with an expanding network of cellular factors [5, 6]. Here we employed SILAC-based interaction screens to independently identify POM121 as a capsid interactor (Fig 1A). POM121 is a transmembrane nuclear pore protein composed of an amino-terminal membrane-anchoring domain and a carboxy-terminal FG-rich region that extend towards the transport channel [28, 29]. POM121 has been recently described as an HIV-1 host factor whose depletion impairs infection [30, 31]. We have built on these results by providing new insights into the breadth and determinants of POM121 dependency during lentiviral infection, and by identifying capsid sequence and associated host factors as modulators of this dependency.

One of the most significant findings of this study is that POM121 depletion suppresses HIV-1 infection in primary CD4+ T cells, the main physiological target of HIV-1 infection *in vivo* (Fig 2E). NPC composition and NUPs requirement can vary substantially amongst cellular backgrounds [35, 37], and recent work has shown that HIV-1 can remodel NPCs in primary CD4+ T cells to facilitate infection [47]. These results therefore confirmed that the requirement for POM121 is not restricted to transformed laboratory cell lines. Consistent with this, POM121 depletion impaired HIV-1 infection across several cell lines, although the magnitude of the phenotype varied depending on the cellular background (Fig 1B). HEK293T cells showed the weakest effect, with an approximately 1.6-fold reduction in infection. Interestingly, Guo and colleagues reported an approximately twofold reduction in HIV-1 infection following siRNA-mediated depletion of POM121 in the same cell type [30], broadly consistent with the modest phenotype observed here.

We found that POM121 dependency is not restricted to HIV-1, as infection by SIVmac and the chimeric HIV-1(SIVmac CA) virus was also impaired in its absence. In striking contrast, EIAV infection was significantly enhanced following POM121 depletion (Fig 2D). EIAV has been shown to be relatively insensitive to depletion of several NUPs that affect HIV-1 infection, while sharing its dependency on others, including NUP153 [35, 41, 42]. However, to our knowledge, this is the first report showing that depletion of a nucleoporin enhances EIAV infection. These findings indicate that POM121 can exert proviral or inhibitory roles, depending on the viral context.

Similarly, and in agreement with previous reports [31, 43, 44], we found that overexpression of cytoplasmic POM121-ΔN, which lacks the nuclear membrane-anchoring region but retains the FG-rich domain, inhibited a broader range of lentiviruses, including HIV-1, HIV-2, HIV-1 (SIVmac CA), SIVmac, and EIAV (Fig 3E). Notably, several viruses that were insensitive to depletion of endogenous POM121 are nonetheless inhibited by POM121-ΔN. This distinction indicates that functional dependence on endogenous POM121 and sensitivity to its isolated C-terminal region are separable properties.

Lentiviral capsids contain a FG-binding pocket that is exploited by several cellular factors [7]. The timing and cellular location at which these factors engage this pocket may determine the outcome of infection. This is best illustrated by the truncated CPSF6-358 (1-358) which lacks the carboxy-terminal RS-rich region required for efficient nuclear localisation and consequently accumulates in the cytoplasm, where it binds the viral capsid [45]. This premature association interferes with the normal sequence of CA interactions with other factors that recognize the same pocket, including NUP153, thereby inhibiting infection [45, 46]. In contrast, our data show that overexpression of POM121-ΔN does not further reduce HIV-1 infectivity in POM121-null cells (Fig 3F). This lack of additivity suggests that, for HIV-1, POM121-ΔN and depletion of endogenous POM121 affect the same or overlapping steps of infection, or that that the inhibitory activity of the C-terminal fragment depends on the presence of full length POM121. However, the broader antiviral activity of POM121-ΔN argues against these perturbations acting through an identical mechanism, since HIV-2 and EIAV are inhibited by POM121-ΔN despite being insensitive to POM121 depletion. Thus, POM121-ΔN is unlikely to act simply as a dominant-negative form that prevents productive interaction between HIV-1 capsid and endogenous POM121. Instead, its premature cytoplasmic association with lentiviral capsids may interfere more broadly with capsid trafficking or with subsequent host-factor interactions, while converging on a POM121-dependent step.

The viral capsid is the genetic determinant for POM121 sensitivity. HIV-1 capsid mutants altered in the CypA-binding loop (P90A), FG-binding pocket (N74D) or tri-hexamer interface (T210K) were all resistant to depletion of endogenous POM121 (Fig 2C). N74D and T210K were also resistant to POM121-ΔN, whereas P90A showed only modest sensitivity to expression of the truncated protein (Fig 3D). These results differ quantitatively from those reported by Xue and colleagues, who observed attenuated sensitivity of P90A and N74D viruses to POM121 depletion [31]. The lack of P90A dependence observed here is consistent with the strong CypA dependence of the POM121 phenotype (discussed below). In contrast, the behaviour of N74D is consistent with the reduced ability of this mutant to engage FG-rich motifs [45].

We also observed a difference with these authors at the level of N74D capsid binding (Fig 4D). Whereas N74D CANC assemblies did not detectably interact with POM121 in our assays, Xue and colleagues reported efficient binding of N74D capsids to a carboxy-terminal POM121 fragment [31]. The constructs used in the two studies were not identical. Here, we analysed the canonical 1,249-residue POM121 protein and a carboxy-terminal fragment spanning residues 683–1,249, whereas Xue *et al*. used a 984-residue POM121 construct lacking the first 265 residues of the canonical protein and a carboxy-terminal fragment corresponding approximately to residues 704–1,249. Whether differences in the construct employed contribute to the distinct N74D-binding phenotypes is currently under investigation. Another important conclusion from our work is the lack of a simple correlation between capsid binding and functional dependence on POM121. N74D showed both loss of detectable POM121 and POM121-ΔN binding and resistance to POM121 depletion. In contrast, wild type, P90A and T210K capsids all interacted with POM121, yet only wild type HIV-1 was impaired by POM121 depletion (Fig 4). This uncoupling between capsid binding and infection dependency is reminiscent of MX2, which can interact with capsid mutants that nevertheless escape MX2-mediated restriction [48, 49].

The interplay between capsid-binding cellular factors critically influences the outcome of HIV-1 infection. This is well illustrated by CypA, whose early engagement of the incoming capsid modulates subsequent host-factor interactions and their functional consequences. CypA shields the capsid from premature CPSF6 binding and recognition by TRIM5α, competes with Transportin-1 (TNPO1) for engagement of the CypA-binding loop, and contributes to determining viral sensitivity to MX2-mediated restriction [16, 35, 46, 50]. To determine whether CypA similarly influences POM121 dependency, we depleted CypA in POM121-deficient cells. Strikingly, loss of CypA fully restored HIV-1 infectivity to the levels observed in control cells (Fig 5A), identifying CypA as a major determinant of the POM121-dependent phenotype. A similar rescue was observed for SIVmac. Given that SIVmac does not efficiently engage CypA [40], this finding suggests that CypA can influence POM121 dependency independently of canonical direct CypA-capsid binding. This framework may also explain the resistance of the P90A mutant virus: despite retaining the ability to interact with POM121, its inability to bind CypA renders infection independent from endogenous POM121.

The T210K mutant deserves further consideration. Its resistance to POM121 depletion is not determined by loss of POM121 binding, as for N74D, or by inability to engage CypA, as for P90A. During the course of our experiments, we uncovered an inhibitory effect of CypA on T210K infection, with CypA depletion increasing infectivity approximately fourfold (Fig 5C). Thr210 is located at the tri-hexamer interface of the capsid lattice, away from the putative CypA-binding loop. We therefore speculate that CypA might inhibit T210K by modulating the interaction with other required cellular factor/s.

Interestingly, recent findings further connect this region of CA with POM121. Replication of viruses in the presence of an inhibitory C-terminal fragment of POM121 (876-1,249) selected the CA substitutions R132G and G208R, which in combination, rendered the virus resistant [44]. G208R maps close to T210 at the tri-hexamer interface. G208R and T210K have also been selected by viruses grown under the selective pressure imposed by MX2, and both confer resistance to it [34, 51]. This is particularly relevant because the amino-terminal RRR motif of MX2 engages the negatively charged tri-hexamer interface of the assembled capsid [39, 52]. More recently, Melčák and colleagues identified a similar stretch of basic residues at the carboxy-terminal end of POM121 that act as a binding enhancer, increasing the affinity of FG-containing POM121 peptides for the capsid [21]. Consistent with this observation, deletion of the final 14 residues from a C-terminal POM121 fragment has been shown to abolish efficient capsid binding and antiviral activity [44]. Together, these observations raise the possibility that POM121 engages the assembled capsid through a bipartite interaction involving both FG-mediated binding and its extreme C-terminal basic region. In this model, T210K could retain detectable POM121 association through interactions involving the FG-binding pocket while disrupting a second interaction involving the tri-hexamer interface, thereby uncoupling POM121 binding from functional POM121 dependency. Further experiments will be required to test this model directly.

POM121/CPSF6 double-CRISPR cells showed an intermediate phenotype. Although loss of CPSF6 partially restored HIV-1 and SIVmac infectivity in POM121 deficient cells, infection did not recover to the levels seen in the presence of POM121. CPSF6 may contribute to the POM121 phenotype at the level of capsid release from the nuclear pore. CPSF6 has been proposed to competitively displace NUP153 from the capsid at the nuclear basket, thereby facilitating release of the viral complex into the nucleoplasm [26]. In this context, loss of POM121 could alter the normal sequence of FG-mediated interactions across the pore, rendering subsequent CPSF6 engagement non-productive. Depletion of CPSF6 could therefore partially relieve this block by allowing the capsid to access alternative nuclear-entry routes, while the incomplete rescue indicates that CPSF6 is only one component of the POM121-dependent pathway. This interpretation is also compatible with the POM121 resistance of N74D, which cannot efficiently engage CPSF6 and therefore may bypass the CPSF6-dependent branch of nuclear entry.

Given its location at the NPC, POM121 may interfere with the nuclear import of the viral replication complex. Indeed, previous studies have shown that POM121 contributes to the nuclear import of viral VPR [53], and that its depletion reduces the accumulation of viral 2-LTR circles, a commonly used surrogate of HIV-1 nuclear entry [30, 31], a phenotype that we also observed here (Fig 6C). We extended these observations by examining whether POM121 contributes to the nuclear transport of other cargos. Using a panel of GFP-LacZ reporters bearing different cellular and viral NLSs [35], we compared their nuclear accumulation in control and POM121 CRISPR cells. Strikingly, POM121 depletion strongly impaired nuclear accumulation of SV40 large T-antigen and N_MX2_ NLS-tagged GFP-LacZ constructs (Fig 6A, 6B). This result confirms that POM121 is not only a key player on the nuclear import of viral replication complexes but is an important regulator of the nucleocytoplasmic transport of viral and non-viral cargo. The strong effect of POM121 depletion on the N_MX2_ reporter is particularly intriguing. The first 25 residues of MX2 contain the motif responsible for targeting the protein to the nuclear envelope and are also critical for its antiviral activity [38, 48, 51, 52, 54]. Our results therefore raise the possibility that POM121 contributes to the nuclear-envelope targeting or transport pathway used by MX2. This would be consistent with our previous work linking MX2 localisation and antiviral activity to nuclear pore-associated proteins, including NUP214 and TNPO1 [38]. However, whether POM121 is required for the localisation or antiviral function of full-length MX2 remains to be determined.

In conclusion, current models propose that largely intact HIV-1 capsids traverse the NPC through sequential interactions with FG-NUPs, potentially facilitated by an affinity gradient across the pore [11–13, 21]. Together, our findings have shown that POM121 participates in this network of capsid-host interaction at the nuclear pore. They further show that functional dependence on a nucleoporin is determined not simply by capsid binding, but by the broader host-factor context in which that interaction occurs.

## MATERIALS AND METHODS

### Cell culture

Human HEK293T, A549, HeLa and parental U87-MG cells were obtained from the American Tissue Culture Collection (ATCC). All cell lines are cultured in Dulbecco’s modified Eagle’s medium (DMEM) supplemented with 10% fetal bovine serum (FBS), L-glutamine and penicillin-streptomycin at 37°C in a 5% CO_2_ atmosphere.

### siRNA transfection

siRNAs against nuclear pore proteins NUPL1, NUP160, NUP54, NUP205 and POM121, and a scrambled control siRNA were obtained from Dharmacon in the form of a smart pool, containing a mix of 4 individual siRNAs, and therefore maximizing the chances of protein depletion. HeLa, U87-MG, A549 and HEK293T cells were seeded at 50-60% confluency in complete DMEM and transfected with siRNA complexed with Lipofectamine RNAiMAX (Thermo Fisher Scientific) at 1:5 ratio (siRNA:reagent) in Opti-MEM, at a final concentration of 10 nM. Cells were transfected twice, 24 h apart, and challenged with HIV-1/GFP 24 h after the second transfection. Infection was quantified 48 h later by flow cytometry as the percentage of GFP-positive cells. An aliquot of the transfected cells was kept to verify protein depletion by immunoblotting.

### HIV-1 molecular clones and retroviral vectors

The HIV-1 NL4.3/ Nef-IRES-Renilla and NL4.3/ Nef-IRES-GFP reporter viruses have been previously described [55]. GFP-expressing retroviral particles based on murine leukaemia virus (MLV/GFP), equine infectious anaemia virus (EIAV/GFP), simian immunodeficiency virus (SIV/GFP), human immunodeficiency virus type 2 (HIV-2), and HIV-1 (HIV-1/GFP) were used as reporter viruses, together with an HIV-1 chimera bearing the SIVmac capsid [HIV-1(SIV CA)] and HIV-1 bearing wild-type, N74D-, P90A- or T210K-capsid (CA). Particles were produced by transient transfection of HEK293T cells. Cells were seeded in 6-well plates at 8.5 × 10^5^ cells per well in 2 mL of medium. After 48h, and immediately before transfection, the medium was replaced with 2 mL of fresh complete DMEM. Cells were co-transfected with a gag-pol packaging plasmid, a GFP-encoding transfer vector, and a vesicular stomatitis virus glycoprotein (VSV-G) envelope plasmid (pMD.G) at a mass ratio of 1:1:0.25, corresponding to 1.6, 1.6 and 0.4 µg per well, respectively (3.6 µg of total DNA). HIV-1/GFP particles were produced with p8.91 [55], an RRL-based GFP transfer vector [57] and pMD.G [58]. MLV/GFP, EIAV/GFP, SIV/GFP, HIV-2/GFP, and CA-mutant HIV-1/GFP particles were produced under identical conditions using the corresponding packaging and transfer plasmids, as described before [55]. DNA was diluted in 200 µL of Opti-MEM, with PEI Max (24765-1, polysciences) at a ratio of 4 µL PEI per µg of DNA (14.4 µL per well). The mixture was vortexed and incubated for 20 min at room temperature before being added to the cells. The culture medium was replaced with pre-warmed fresh medium ∼16–18 h post-transfection. Viral supernatants were harvested 48 h post-transfection, clarified through 0.45-µm syringe filters, dispensed into 1 mL aliquots and stored at −80 °C until use. Viral titers were determined by infecting 3 x 10^4^ HeLa cells with serial dilutions of viral supernatants and quantifying GFP-positive cells by flow cytometry 48 h post-infection.

### CRISPR-Cas9 cell lines generation

POM121, cyclophilin A (CypA) and CPSF6 were depleted by CRISPR-Cas9 genome editing using the lentiCRISPRv2 Puro, Hygro or Blast system, respectively [59]. Guide RNAs were cloned into BsmBI-linearised lentiCRISPRv2 using annealed oligonucleotides. The 20-nt protospacer sequences (5ʹ→3ʹ) were CTCCTCCCGCTCCCAGACAC for POM121, GCCCGACCTCAAAGGAGACG for CypA, and CATGTAATCTCGGTCTTCTG for CPSF6. Control cells were transduced with a vector expressing a non-targeting sgRNA (GATATGTGTGATGAGCTCGC).

Lentiviral particles were produced in HEK293T cells by transfection of the packaging plasmid p8.91, pMD.G, and the corresponding lentiCRISPRv2 construct at a ratio of 1:0.25:1. At 48 h post-transfection, supernatants were clarified and used to transduce HeLa cells, which were selected with 20µg/mL puromycin (InvivoGen) for POM121 CRISPR cells, with 200µg/mL hygromycin (InvivoGen) for CypA CRISPR cells, and with 25µg/mL blasticidin (InvivoGen) for CPSF6 CRISPR cells. Single-cell clones were derived from the POM121 bulk population by limiting dilution in 96-well plates. Loss of POM121, CypA or CPSF6 was confirmed by immunoblotting. To generate double-CRISPR lines (POM121/CypA and POM121/CPSF6), POM121 CRISPR cells were additionally transduced with the corresponding guides.

### Analysis of primary human CD4+ T cells

Human primary CD4+ T cells were obtained from peripheral blood mononuclear cells (PBMCs) from healthy donors (approved by the Guy’s Research Ethics Committee) (ref. 03/02/06), and through the Infectious Diseases BioBank at King’s College London (ethics reference MM2-220518) under overall permission from the Southampton and South West Hampshire Research Ethics Committee (B) (REC reference 19/SC/0232). CD4+ T cells were isolated from PBMCs using LymphoPrep (Axis-Shield) and the CD4+ T Cell Isolation Kit (Miltenyi Biotec). Anti-CD4 (BD Pharmingen, 555349) and anti-CD3 (BD Pharmingen, 555340) antibodies were used to determine the purity of isolated cell populations. Values over 90%, as determined by flow cytometric analysis, were consistently obtained. Cell activation was achieved by addition of Dynabeads Human T-Activator CD3/CD28 (Thermo Fisher) and 50 U ml^−1^ recombinant IL-2 (rIL-2) (Roche) for 48 h in RPMI 1640-GlutaMax medium supplemented with 10% heat-inactivated autologous serum, penicillin (100 U ml^−1^) and streptomycin (100 μg ml−1). After activation, CD4+ T cells were maintained in same medium containing 30 U ml^−1^ rIL-2.

Cells were transduced with short hairpin RNAs (shRNAs) targeting POM121 (guide 1: gatccGCAGCAGTGTGAGTACCACTTCAAGAGAGTGGTACTCACACTGCTGCTTTTTTg, guide 2: gatccGCAGCAGCTCGGTGTTTGGTTCAAGAGACCAAACACCGAGCTGCTGCTTTTTTg, guide 3: gatccGTTGACAGGCGCTTACGCAATTCAAGAGATTGCGTAAGCGCCTGTCAATTTTTTg) or a scrambled-sequence control shRNA [60], as described before [55]. Following transduction, cells were challenged with the HIV-1 NL4.3/Nef-IRES-Renilla reporter virus and infection was measured 48 h later as Renilla luciferase activity. POM121 depletion was evaluated by immunoblotting, and infection measurements were compared between POM121-targeting and control shRNA conditions for each donor.

### Plasmids

cDNAs encoding Flag-tagged full-length POM121, POM121-ΔC (amino acids 1–682), POM121-ΔN (amino acids 683–1249) (residue numbering refers to the canonical 1,249-amino-acid protein (UniProt Q96HA1)), or MX1 (negative control) were inserted into a lentiviral-based transfer vector expressing the puromycin resistance gene (IRES-puromycin N-acetyl-transferase, puromycinR) by standard cloning techniques. Lentiviral particles encoding puromycinR plasmids were produced in HEK293T cells as described above, using p8.91 [56], pMD.G [58] and the corresponding transfer plasmid. HeLa cells were seeded at 1.6 × 10⁵ cells per well in 6-well plates and transduced with the filtered supernatant. The medium was replaced 6–8 h later, puromycin added at 1 µg/mL, and cells selected for 72 h. Expression of each protein was verified by immunoblotting.

MLV Capsid-Nucleocapsid (CANC) cDNA (residues 215-534 from MLV Pr65 Gag) were inserted into pST39 [61] using the NdeI and XhoI sites. The HIV-1 CANC construct has been previously described [48].

### Infection assays

3 x 10^4^ cells were seeded in 96-well-plates and challenged with HIV-1/GFP, the CA-mutant HIV-1/GFP viruses, or the HIV-2, HIV-1(SIV CA), SIVmac, EIAV or MLV reporter viruses. The percentage of GFP-positive cells was determined 48 h later by flow cytometry. All infections were repeated at least three times.

### Stable isotope labelling by amino acids

293T cells were grown in reduced DMEM media with no glutamine, no lysine, and no arginine (Thermo Fisher), and supplemented with 10% dialysed fetal bovine serum, 6 mM l-glutamine, 100 U/ml penicillin, 100 μg/ml streptomycin, containing either: (1) 73 mg/l ^13^C_6_,1^5^N_2_-L-lysine (Lys8) and 42 mg/l ^13^C_6_,^15^N_4_-L-arginine (Arg10) (CK Isotopes) for ‘heavy’ conditions; (2) 73 mg/l 4,4,5,5-D4-L-lysine (Lys4) and 42 mg/l ^13^C_6_-L-arginine HCl (Arg6) (CK Isotopes) for ‘medium’ conditions; or (3) 73 mg/l L-lysine (Lys0) and 42 mg/l L-arginine (Arg0) (Sigma) for ‘light’ conditions. Cells were grown on these media for at least 14 days prior to their use.

### CANC Pull-down assays

CANC proteins (wild-type, P90A, T210K or N74D CANC complexes) were expressed in *Escherichia coli* Rosetta (DE3) cells (Merck Millipore) as previously described [32]. Assembled CANC complexes were prepared by dilution of the protein to 40 μM final concentration in 50 mM Tris-HCl pH 8 containing 100 mM NaCl and adding 5 μM of TG50 oligonucleotide (5ʹ-TG(25)-3ʹ). The reaction was incubated overnight at room temperature. For pull-down experiments, protein overexpression was achieved by transient transfection of 293T cells with vectors expressing Flag-tagged proteins (Flag-MX1, Flag-CPSF6, Flag-POM121, Flag-POM121ΔN and Flag-POM121ΔC). At 48 h after transfection, cells were lysed in hypotonic lysis buffer (10 mM Tris-HCl pH 8, 10 mM KCl, 1×complete protease inhibitor cocktail (Roche)) using a Dounce homogenizer. Cell lysates were cleared by centrifugation at 20,000 g for 15 min at 4 °C. Two-hundred microliters of the lysate were mixed with either 40 μl of CANC assembly buffer (50 mM Tris-HCl pH 8, 100 mM NaCl) containing 5 μM TG50 (as a control), or with 40 μl of 40 μM assembled CANC; an input sample was withdrawn from the latter. The mixtures were incubated for 1 h at room temperature, then overlaid on 250 μl of 70% sucrose cushion. Following centrifugation at 20,000 g for 15 min at room temperature, a sample of the resulting supernatant was withdrawn, and the pellet was washed with 500 μl of wash buffer (50 mM Tris-HCl pH 8, 50 mM NaCl and 5 mM KCl), and re-pelleted at 10,000 g for 8 min. Finally, the washed pellet was resuspended in 50 μl sample buffer. Input, supernatant and pellet fractions were analysed by immunoblotting using anti-CA [62] and anti-Flag (Sigma, A8592) antibodies. All experiments were repeated at least three times.

### HIV-1 CANC-interacting proteins and mass spectrometry

SILAC-labelled lysates were prepared from U87-MG cells grown in ‘light’, ‘medium’ or ‘heavy’ media (groups of 4×10 cm dishes per labelling condition) by disrupting the cells in 4 ml of lysis buffer (PBS, 0.5% Triton), and clearing the resulting lysate by centrifugation at 20,000 g, 15’, at 4°C. Capsid-nucleocapsid (CANC) assemblies of wild-type HIV-1, mutant P90A, and MLV were prepared as described under ‘CANC pull-down assays’, and CANC assembly buffer containing 5 µM TG50 was used as a mock control. The following side-to-side experiments were performed: wild type CANCs (light) *versus* MLV CANCs (heavy) (n = 2 biological replicates), P90A CANCs (medium) *versus* MLV CANCs (heavy) (n = 2), wild type CANCs (medium) *versus* mock (heavy) (n = 2) and P90A CANCs (light) *versus* mock (heavy) (n = 2).

In these experiments 250 µl of 40 µM assembled CANC or 250 µl of CANC assembly buffer (50 mM Tris-HCl pH 8, 100 mM NaCl) containing 5 μM TG50 were incubated with the corresponding lysate for 1 h at room temperature. CANC mixtures were then pulled down by centrifugation through a 20% sucrose cushion at 20,000 g for 15’ at 21°C. Recovered pellets were washed with wash buffer (50 mM Tris-HCl pH 8, 50 mM NaCl and 5 mM KCl) and centrifuged again at 10,000 g for 8 min at 21°C. Finally, the pellets were resuspended in sample buffer, pooled, and resolved using 12% SDS–polyacrylamide gels and stained with SimplyBlue (Thermo Fisher).

Proteomic analysis of the resulting samples was performed as previously described (Betancor *et al.*, 2021). Briefly, in-gel reduction, alkylation and tryptic digestion were performed, followed by peptide extraction through successive washes with acetonitrile and water. Extracted peptides were lyophilised, resuspended in 50 mM NH4HCO3 and subjected to LC-MS/MS analysis. Chromatographic separation was performed on an EASY nLC-II nanoflow system (Thermo Fisher) using reversed-phase chromatography with a 75 μm C18 PepMap column. Peptides were separated using a three-step linear gradient of acetonitrile in 0.1% formic acid at a flow rate of 300 nL/min. The resulting eluate was ionised by electrospray ionisation and analysed using an Orbitrap Velos Pro mass spectrometer (Thermo Fisher) controlled by Xcalibur v2.2 software. Data acquisition was conducted in automated data-dependent mode, whereby precursor ions were selected according to their intensity and subjected to collision-induced fragmentation using a Top-20 collision-induced dissociation method. Collision energy profiles for MS/MS analysis were determined according to the mass-to-charge ratio (m/z) and charge state of each peptide.

Protein identification was performed using MaxQuant v1.4.3.14 (Max Planck Institute of Biochemistry) by searching against the UniProt human proteome database (https://www.uniprot.org/proteomes/UP000005640). Search parameters included carbamidomethylation of cysteine residues as a fixed modification, while N-terminal acetylation and methionine oxidation were considered variable modifications. A minimum peptide length of seven amino acids and a maximum of five modifications per peptide were permitted. Peptide-spectrum matches and protein identifications were filtered using a false discovery rate threshold of 1%. Subsequent data processing was carried out with Perseus software (v1.4.1.3), where potential contaminants and reverse-sequence protein identifications were excluded before determining the relative abundance of identified proteins across samples. Representative pairwise comparisons are presented in Fig 1a. The mass spectrometry proteomics data have been deposited to the ProteomeXchange Consortium via the MassIVE partner repository with the dataset identifier PXD086878 (MSV000103847).

### Immunoblotting

Cells were lysed in buffer containing 10 mM Tris-HCl pH 8, 150 mM NaCl, 1 mM EDTA, 1% Triton X-100 and 0.1% sodium deoxycholate. Lysates were briefly sonicated, boiled for 10 min and resolved in Mini-PROTEAN TGX precast gels (Bio-Rad). After electrophoretically transferring proteins to a 0.45 μm nitrocellulose membrane (Amersham), their presence was analysed using specific antibodies.

### Immunofluorescence microscopy

HeLa cells (CRISPR control, POM121 CRISPR clones, or cells stably expressing full-length POM121, POM121-ΔN or POM121-ΔC) were seeded onto coverslips at ∼20,000 cells per well in 24-well plates. At 24 h, cells were washed three times with 1× phosphate buffer saline and fixed in 4% paraformaldehyde (EM Sciences) for 15 min before permeabilization with 0.2% Triton X-100 for 15 min. The specimens were blocked in buffer NGB (50 mM NH4Cl, 2% goat serum and 2% bovine serum albumin) for 1 h.

Endogenous NUPs were detected using mouse anti-NUPs monoclonal antibody (Sigma, N8786) and secondary donkey anti-mouse antibody conjugated to Alexa 594 (Invitrogen, A-21203). Endogenous POM121 was detected using rabbit anti-POM121 polyclonal antibody (proteintech, 15645-1-AP) and secondary donkey anti-rabbit antibody conjugated to Alexa 488 (Invitrogen, A-21206). DAPI staining was used to demarcate the nucleus (0.1 mg/ml for 5 min).

In experiments directed at determining the effect of HIV-1 infection in POM121 cellular distribution, HeLa cells expressing Flag-tagged full-length POM121, POM121-ΔN or POM121-ΔC were challenged with HIV-1/GFP 16 h prior fixation. Flag-tagged proteins were detected using a mouse anti-Flag monoclonal M2 (Sigma, F3165) and secondary donkey anti-mouse antibody conjugated to Alexa 594 (Invitrogen, A-21203). DAPI staining was used to demarcate the nucleus (0.1 mg/ml for 5 min).

Nuclear import was assessed using a panel of nine chimeric GFP-LacZ reporters fused to different nuclear localisation signals (NLSs), as previously described [35]. In brief, retroviral LHCX-based vectors containing different NLS-GFP-LacZ constructs were used to transduce HeLa cells. At 48 h, 600 μg ml^−1^ hygromycin (Invitrogen) was added for 5 days and hygromycin-resistant cells were seeded on coverslips and stained with DAPI.

In all cases, cells were visualized using a Zeiss LSM 800 confocal microscope with Airyscan. Nuclear accumulation was quantified per cell as the Manders’ colocalisation coefficient M1, defined as the fraction of GFP signal overlapping the DAPI-defined nuclear mask, calculated with the JACoP plugin in Fiji [63].

### Quantification of HIV-1 2-LTR

Control HeLa cells (CRISPR CTR) and HeLa cells with CRISPR-mediated knockout of POM121 (CRISPR POM121 c1 and c2) were infected with VSV-G-pseudotyped HIV-1 NL4.3/Nef-IRES-GFP reporter virus. Non-infected HeLa cells were also collected to use as a negative control. Cells were resuspended in 100 µL of PBS and lysed in 500 µL of genomic DNA extraction buffer (100 mM Tris-HCl, pH 8.0, 5 mM EDTA, 0.2% SDS, and 50 mM NaCl). Two microliters of proteinase K (20 mg/mL) and RNase A (10 mg/mL) were added, and samples were incubated at 50 °C for 1 h. DNA was extracted using phenol/chloroform/isoamyl alcohol, followed by phase separation using Phase Lock Gel tubes. The aqueous phase was collected, and DNA was precipitated with isopropanol and 3 M sodium acetate (pH 5.4). The DNA pellet was washed twice with 70% ethanol, air-dried, and resuspended in 50 µL of water. The extracted DNA was used as a template for the quantification of HIV-1 2-LTR circles by quantitative PCR (qPCR).

To generate a standard, a fragment spanning the 2-LTR junction was amplified from an infected sample with Taq DNA polymerase using primers 5ʹ-AACTAGGGAACCCACTGCTTAAG-3ʹ (forward), and 5ʹ-TCTTGTCTTCTTTGGGAGTGAATT-3ʹ (reverse). The resulting PCR product was cloned into a TOPO vector using the TOPO™ TA Cloning Kit (ThermoFisher, K4575J10), following the manufacturer’s instructions. The recombinant plasmid was verified by DNA sequencing to confirm the identity and sequence of the cloned 2-LTR junction. The verified plasmid was subsequently used to prepare a serial dilution standard curve for qPCR efficiency validation.

2-LTR circles were quantified by SybrGreen qPCR (2X Universal SYBR Green Fast qPCR Mix, RK21203, ABclonal) using 10 ng of extracted DNA from infected cells and the same 2-LTR primers (final concentration 1µM). To account for differences in the amount of cellular DNA between samples, amplification of the cellular reference gene *TRIM5* was used for normalization. The primers for *TRIM5* were 5ʹ-GCTCACCAAGCTCTTGAACCTT-3ʹ (forward) and 5ʹ-GTGTATGAGATGCACCTGGACA-3ʹ (reverse). The qPCR conditions included an initial denaturation of 95°C for 10 min, followed by 40 cycles of denaturation (95°C for15 s) and annealing/extension (60°C for 1 min).

2-LTR fold change was calculated after normalization to the cellular reference gene and expressed relative to the control HeLa cells. The equations used were:

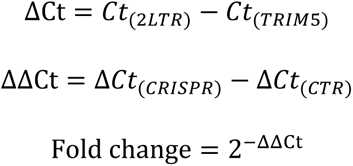

### Aphidicolin treatment

To block cell cycle progression and DNA replication, CRISPR POM121 clones were treated with aphidicolin (2 µg/ml) or an equivalent volume of vehicle (DMSO) for 24 h prior to infection with HIV-1/GFP and maintained in the same media for the course of the infection. The percentage of GFP⁺ cells was determined by flow cytometry as described above. Fold inhibition was calculated as the ratio of the percentage of GFP⁺ cells in DMSO-treated cells in comparison to aphidicolin-treated cells.

### Statistical analysis

Data are presented as the mean ± standard deviation (SD), unless otherwise indicated. Statistical comparisons between experimental conditions and their respective controls were performed using unpaired Student’s *t*-test, as indicated in the figure legends. The number of biological replicates and the statistical significance threshold were specified for each experiment. A *p* value below 0.05 was considered statistically significant.

## ACKNOWLEDGMENTS

This work was supported by the Spanish Ministry of Science and Innovation and the Agencia Estatal de Investigación (MCIN/AEI/10.13039/501100011033) through a Ramón y Cajal fellowship (RYC2021-032674-I), funded by the European Union NextGenerationEU/PRTR, and a Proyectos de Generación de Conocimiento grant (PID2022-140647NA-100) co-funded by the European Regional Development Fund (ERDF), European Union, and by the Wellcome Trust (106223/Z/14/Z and 222433/Z/21/Z to MHM). Laura Arribas was supported by a predoctoral FPI fellowship associated with grant PID2022-140647NA-100 (PREP2022-000392).

We thank the Advanced Confocal and Electron Microscopy Research Service (SIMACE) at the Universidad de Las Palmas de Gran Canaria (ULPGC) for providing access to microscopy facilities and technical support.

We thank Stelios Papaioannou and Antonio Tugores for the provision of reagents and helpful discussions.

