## Supplementary Figures 1-7 for "POM121 regulates lentiviral nuclear import in a capsid-host factor-dependent manner"

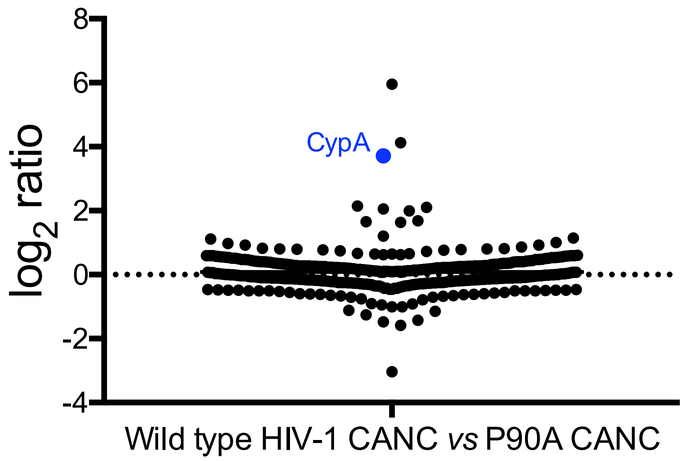


**S1 Fig. SILAC screen comparison between wild type HIV-1 CANC and P90A CANC.** Wild type and mutant P90A HIV-1 CANC assemblies were incubated with HEK293T lysates from cells labelled with 'light', 'medium' or 'heavy' amino acids. Mixtures were subjected to centrifugation through a 70% sucrose cushion, washed with wash buffer, centrifuged again, and analysed by tandem mass spectrometry. CypA is highlighted in blue.

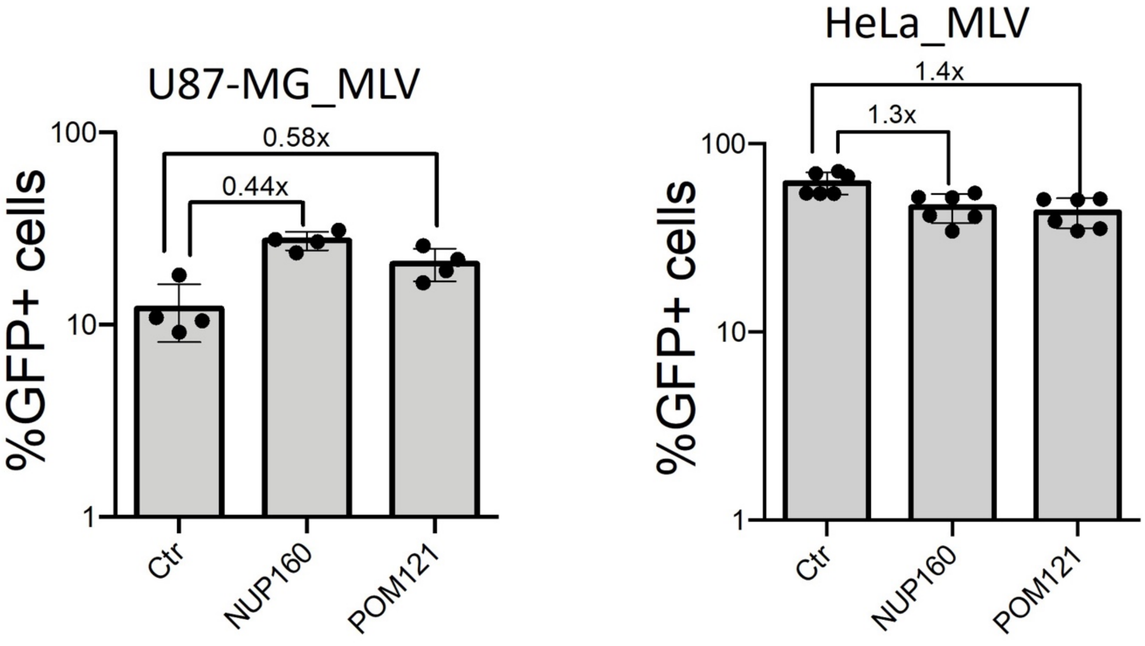


**S2 Fig. MLV is not inhibited by NUPs depletion.** HeLa and U87-MG cells were transfected twice, 24 h apart, with 20 nM of siRNAs against *POM121* and *NUP160*, with a non-targeting siRNA included as a control, Ctr). 24 h later, cells were challenged with a MLV based lentiviral vector expressing GFP (MLV/GFP) and the percentage of infected (GFP-expressing) cells quantitated 48 h later by flow cytometry. The fold reduction in viral infectivity between cells transfected with Ctr siRNA and those transfected with siRNAs against *NUP160* and *POM121* is shown (n = at least 4; mean ± standard deviation (SD); * p-value < 0.05; unpaired t-test to Ctr siRNA).

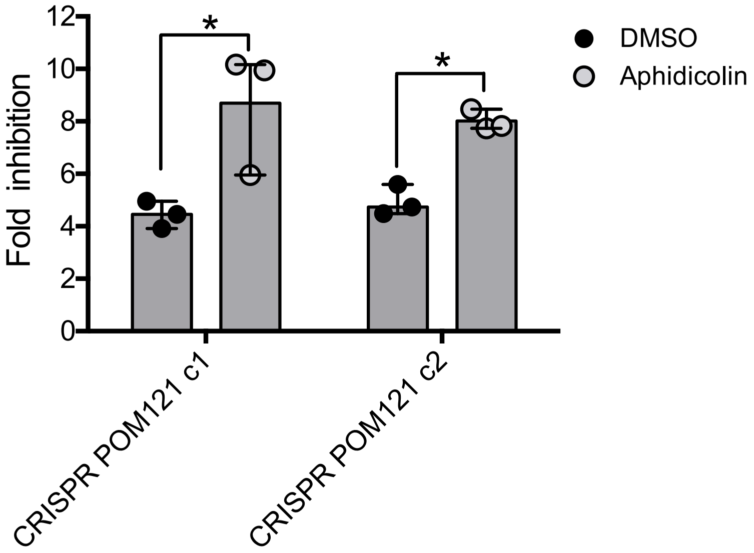


**S3 Fig. Inhibition of cell division enhances the HIV-1 dependence on POM121.** HeLa POM121 CRISPR cells (clones 1 and 2) were treated with 2 µg/ml of aphidicolin, or the same concentration of vehicle DMSO for 24 h, and then challenged with HIV-1/GFP. Aphidicolin/DMSO treatment was maintained during the 48 h course of the infection, and cells were fixed and the percentage of infected (GFP-expressing) cells quantitated by flow cytometry. The fold reduction in viral infectivity between cells treated with DMSO and aphidicolin is shown (n = 3; mean ± standard deviation (SD); * p-value < 0.05; unpaired t-test to Ctr siRNA).

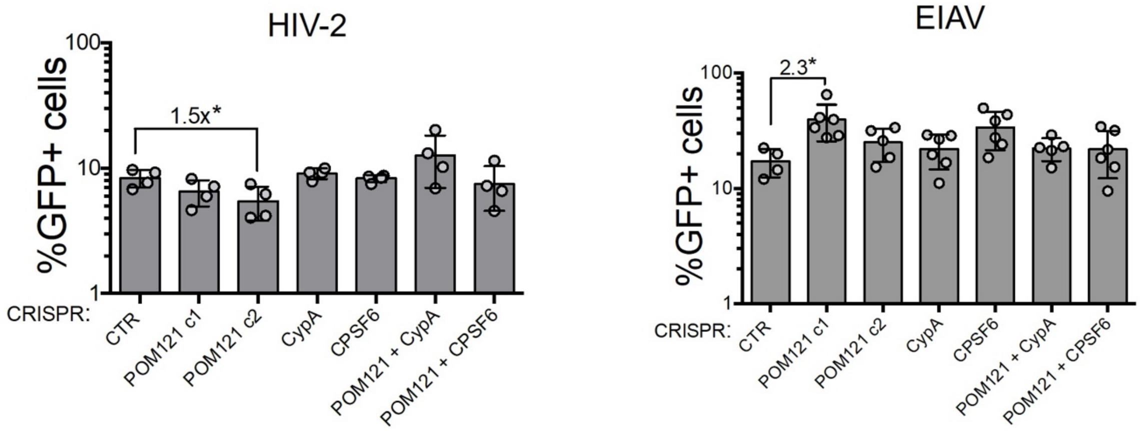


**S4 Fig. Effect of CypA and CPSF6 depletion in POM121 dependence.** Parental HeLa cells and HeLa POM121 CRISPR cells were depleted of CypA or CPSF6 by CRISPR-Cas9 genome editing to generate CRISPR CypA, CRISPR CPSF6, CRISPR POM121 + CypA, and CRISPR POM121 + CPSF6 cells. These cells, together with CTR CRISPR cells, were challenged with HIV-2/GFP (left) and EIAV/GFP (right) particles (left) and infectivity quantitated 48 h later by flow cytometry as the number of GFP+ cells (n = at least 4 biological replicates; mean ± SD; * p-value < 0.05; unpaired t-test to CTR cells).

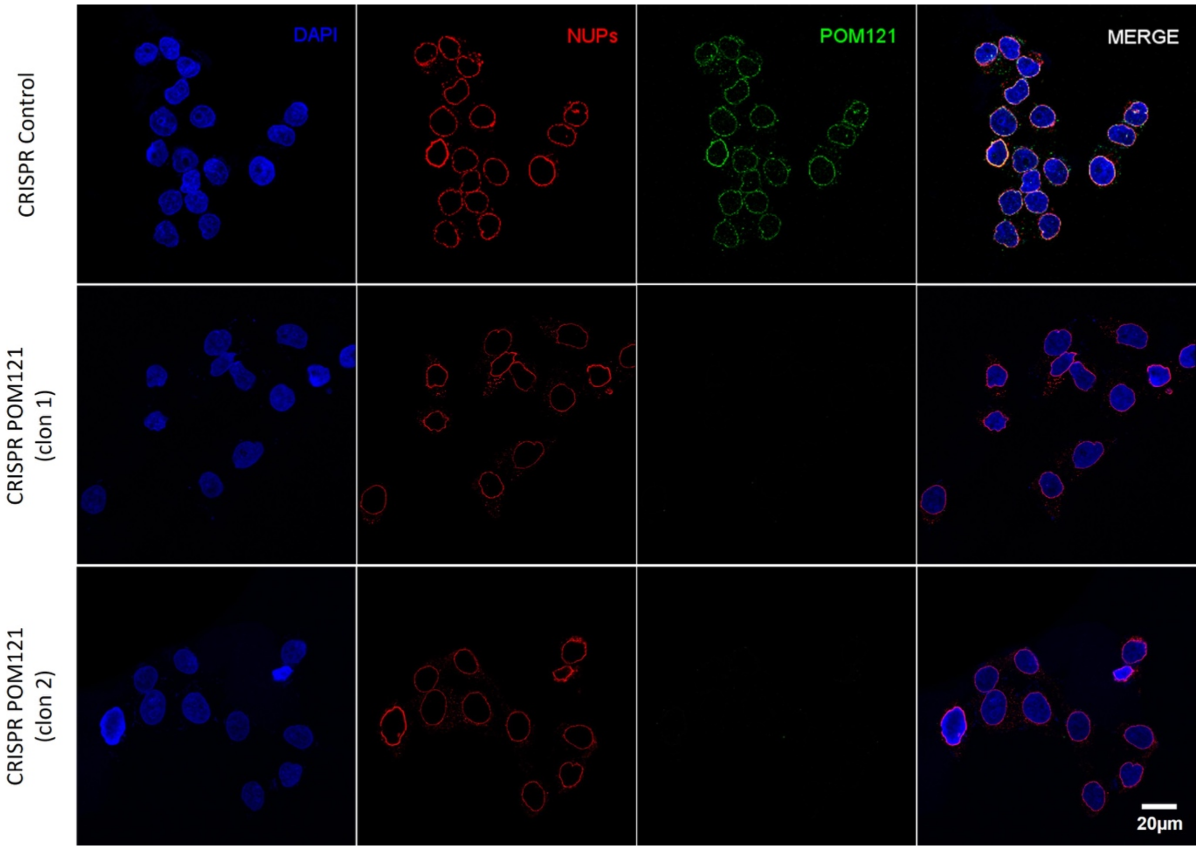


**S5 Fig. POM121 depletion by confocal microscopy.** Hela control CRIPSR and HeLa POM121 CRISPR clones 1 and 2 cell lines were seeded onto glass coverslips, fixed, and permeabilized. POM121 was detected using primary anti-POM121 antibodies and secondary anti-rabbit$^{488}$ antibodies. Mab414 antibody was used to detect FG NUPs, coupled with anti-mouse$^{594}$, and the nuclei was stained with DAPI.

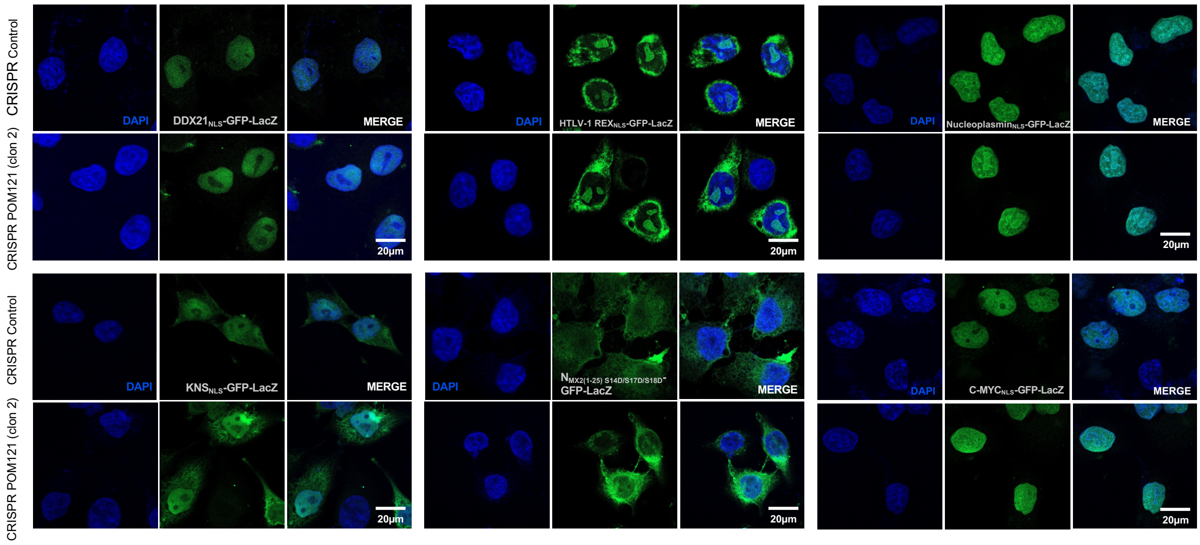


**S6 Fig. Effect of POM121 depletion on the nuclear import of viral and cellular cargo.** HeLa control CRISPR and CRISPR POM121 cells stably expressing chimeric proteins $DDX21_{NLS}$-GFP-LacZ, HTLV1-$REX_{NLS}$-GFP-LacZ, $Nucleoplasmin_{NLS}$-GFP-LacZ, $KNS_{NLS}$-GFP-LacZ, C-$MYC_{NLS}$-GFP-LacZ and $N_{MX2(1-25)\ S14D/S17D/S18D}$-GFP-LacZ (negative control) were seeded on glass coverslips. Cellular distribution of chimeric proteins was detected by confocal microscopy, with the nuclei stained with DAPI.

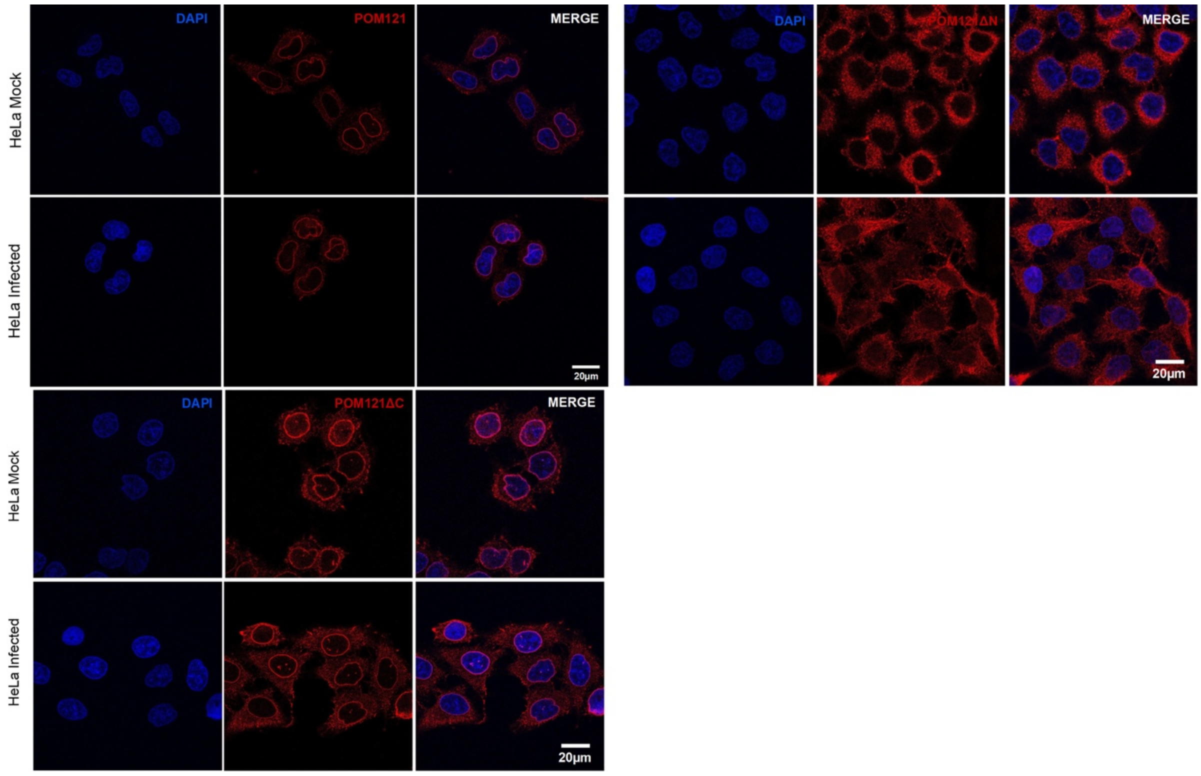


**S7 Fig. Effect of HIV-1 infection in POM121-ΔN and POM121-ΔC cellular distribution.** Hela cells were stably transduced with puromycinR lentiviral vectors encosing with Flag-tagged POM121, POM121-ΔN or POM121-ΔC. Cells were split into two wells and seeded onto glass coverslips, with one of the wells being infected with HIV-1/GFP (MOI = 1), and the other left uninfected. Sixteen h after infection cells were fixed and POM121 forms detected by confocal microscopy using an anti-flag antibody (M2). DAPI was used to stain the nuclei.
